# Towards estimating the number of strains that make up a natural bacterial population

**DOI:** 10.1101/2023.02.20.529252

**Authors:** Tomeu Viver, Roth E. Conrad, Luis M. Rodriguez-R, Ana S. Ramírez, Stephanus N. Venter, Jairo Rocha-Cárdenas, Mercè Llabrés, Rudolf Amann, Konstantinos T. Konstantinidis, Ramon Rossello-Mora

## Abstract

What a strain is and how many strains make up a natural bacterial population remain elusive concepts despite their apparent importance for assessing the role of intra-population diversity in disease emergence or response to environmental perturbations. To advance these concepts, we sequenced 138 randomly selected *Salinibacter ruber* isolates from two solar salterns and assessed these genomes against companion short-read metagenomes from the same samples. The distribution of genome-aggregate average nucleotide identity (ANI) values among these isolates revealed a bimodal distribution, with significantly lower occurrence of values between 99.2% and 99.8% relative to ANI >99.8% or <99.2%, revealing a natural “gap” in the sequence space within species. Accordingly, we used this ANI gap to define genomovars and a higher ANI value of >99.99% and shared gene-content >99.0% to define strains. Using these thresholds and extrapolating from how many metagenomic reads each genomovar uniquely recruited, we estimated that –although our 138 isolates represented about 80% of the *Sal. ruber* population– the total population in one pond is composed of 5,500 to 11,000 genomovars, the great majority of which appear to be rare *in situ*. These data also revealed that the most frequently recovered isolate in lab media was often not the most abundant genomovar *in situ*, suggesting that cultivation biases are significant, even in cases that cultivation procedures are thought to be robust. Preliminary analyses of available genomes revealed that the thresholds used for defining strains and distinct intra-species units (genomovars) may be broadly applicable to additional bacterial species.

**Significance Statement:** Strains are the smallest distinguishable units within a microbial species. Strains that carry unique gene content often underly the emergence of disease outbreaks and the response of the species to environmental perturbations. Therefore, a major challenge in microbiome research across environmental and clinical settings is to evaluate how many strains of the same species coexist in nature and how dominant strains emerge from this diversity. Unfortunately, the available theoretical concept of strain is not directly applicable to culture-independent surveys. Here, we provide such a practical definition for strain and use it to show that that the number of strains making up a natural bacterial population may be large, in the order of a few thousands, but not infinite.

## Introduction

A prokaryotic species is composed of by multiple strains, the smallest distinguishable units within species, which typically show higher than 95% ANI among themselves vs. <90% ANI to strains of different species (1, 2). Strains that show intermediate identities (e.g., 90-95% ANI) are comparatively much less frequent, or even absent for some taxa, relative to strains showing >95% ANI under the same environmental conditions and thus, the former strains presumably represent distinct species (3, 4), revealing a natural gap or discontinuity in genome diversity at the species-level. The strains of a species frequently carry a high degree of inter-strain sequence (allelic) diversity in core genes and strain-specific auxiliary gene content (5, 6). This diversity often underlies the emergence of disease outbreaks (7, 8) and/or the response of the species to environmental perturbations (9). Therefore, a major challenge in microbiome research across environmental and clinical settings is to evaluate how many strains of the same species coexist in the environment in order to better quantify intra-species diversity and understand how dominant strains emerge from this diversity. This brings up another related challenge: how to precisely define strains.

Bacteriologists have an operational concept of strain, which has its basis on the pure culture approach, and considers a strain as “*a group of genetically similar descendants of a single colony or cell*” (10). Therefore, strain embraces all derivative lines of a single isolate, regardless of whether or not the descendants have undergone mutational events such as gene loss, duplications, genomic rearrangements, or modifications of the gene expression, as long as these do not affect the key (known) phenotypic properties of the strain. However, this concept is ambiguous because phenotypic similarity often depends on the growth conditions. For example, the isolation of an organism (wild-type) in the laboratory is commonly accompanied by changes in at least gene expression (11), and often gene mutations or deletions (12, 13), due to adaptation to the laboratory conditions. Some of these changes could lead to substantial phenotypic differences; yet, the wild-type and the lab-adapted cells are typically considered the same strain (14). In surveys of natural populations, where strain ancestry information is typically unavailable, strains have been discerned instead based on single nucleotide variants patterns (SNVs), but even in such cases a widely accepted definition on the number of SNVs expected to define a strain has not emerged yet (8, 15). Note also that strain should not be equated to ‘clone’ because the latter implies identical genetic sequence at selected loci or/and the whole-genome (16, 17), which is not a prerequisite for members of the same strain.

To advance the current definition of strain, we used a large collection of isolates of the model hypersaline species *Salinibacter ruber* from two solar saltern sites in the Mallorca and Fuerteventura Islands, Spain (18). Solar salterns are human-controlled tanks or ponds, used for the harvesting of salt for human consumption. These ponds are operated in repeated cycles of feeding with natural saltwater, increasing salt concentration due to water evaporation, and finally, salt precipitation. Several studies have shown that salterns in different parts of the world harbour recurrent microbial communities each year, characterised by low diversity of higher taxa (family-level and above), generally consisting of two major lineages i.e., the archaeal *Halobacteria* class and the bacterial class *Rhodothermia* (19), but with relatively high genus and species richness within each class (9, 20, 21). Importantly, our previous studies have shown that *Sal. ruber* typically makes up about 5-10% of the total microbial community in most saltern sites across the world, including the two salterns sampled herein. Further, these abundant *Sal. ruber* populations typically harbor a large number of distinct genotypes (11, 22), comparable to the diversity of genotypes of the model bacterial species, *Escherichia coli*, that can be found in the public databases and have been isolated from different sources (22). Hence, *Sal. ruber* represents an ideal environmental group of bacteria to study intra-species differentiation and units (18). Here, the genome sequences of 138 *Sal. ruber* isolates, randomly chosen from our larger isolate, and the quantification of their diversity and *in situ* abundance patterns using the metagenomes of the saltern of origin allowed us to propose a natural definition for a “strain” and other sub-species categories, as well as to evaluate the number of strains co-existing in their hypersaline ecosystem.

## Results

### Collection of isolates and identification of their clonal varieties (CVs) using PCR-amplicon fingerprint profiles (RAPD)

The isolates used in this study were recovered from two locations in Spain. The first collection was obtained from four adjacent ponds, fed with the same source seawater, in the ‘Es Trenc’ solar salterns on the Island of Mallorca during perturbation experiments performed over a period of one month in 2012. These experiments manipulated sunlight intensity through the application of shading mesh on the top of the ponds and salinity level through dilution with freshwater (9, 23). In total, 409 randomly picked pure cultures were isolated on standard growth media for *Salinibacter* (19, 22), 207 of which were tentatively identified as *Salinibacter ruber* based on MALDI-TOF mass spectroscopy profiles (22). A cost-effective and widely used approach to discern between putatively identical isolates (clones) is the random amplified polymorphic DNA (RAPD) PCR assay (24). The isolates were dereplicated into clonal varieties (CVs) using up to three different RAPD primer sets. We considered two isolates showing identical RAPD patterns as being the same clonal variety (CV), although identical fingerprints do not necessarily guarantee identical genomes (24). The 207 isolates represented 187 distinct CVs. Of them, 118 randomly selected isolates, representing 107 CVs, including isolates from the same CV, were subjected to whole-genome sequencing. Therefore, these isolates represented both inter- (n=103) vs intra-CV (n=15) diversity as well as the four ponds from the previous treatments (i.e., control, long- and short-shaded, and diluted; n= ~30 isolates/treatment) (22). Specifically, the 15 isolates represented four CVs with two or more representatives (Table 1 and Fig. S1), whereas the remaining 103 had unique profiles (single-isolate CVs). From the other location, the solar salterns on the Fuerteventura Island (Canary Islands), we obtained 46 isolates from a single sample of a salt-saturated (control) pond, 40 of which were identified as *Sal. ruber*. RAPD signatures identified 26 different CVs, and 25 isolates (56%) showed a unique CV whereas the remaining 15 showed identical RAPD profiles. Nine of the latter isolates were sequenced and found to belong to 2 CVs (CV5 and CV6; see below). In addition, 11 of the non-clonal isolates were sequenced and included in this study (Table 1 and Fig. S1).

**Table 1:** Statistics of the genome sequences used in this study and comparisons between genomes assigned to the same clonal variety (CV) vs. different CVs. Rows that include “unique” in the second column represent comparisons between genomes assigned to different genomovars; numbers within parenthesis in the 3^rd^ column represent the number of comparisons performed by randomly splitting the reads of a genome sequencing dataset into two halves (subsampling) and comparing the two halves (i.e., rows representing within genome comparisons); the remaining rows represent comparison between genomes assigned to the same genomovar. Samples CM, DZ, and UM yielded only isolates that each was assigned to a different CV and were not used for the subsampling calculations. Samples and isolates recovered from the Mallorca solar salterns are named according to the different ponds: control (C), dilution (D), and unshaded (U) at three time points: time-zero (Z), 1-week (W), and 1-month (M) (e.g., sample “UM” denotes Unshaded at 1 month) (22). The sample and isolates from Fuerteventura area denoted by “FV” in the name.

| Sample | CVs | Isolates | Nr. genomes<br>(Nr. Subsampled genomes) | Genome size |  | Num. CDs |  | G+C content |  | ANI |  | % Genomic fraction shared |  | Num. SNPs |  |
| --- | --- | --- | --- | --- | --- | --- | --- | --- | --- | --- | --- | --- | --- | --- | --- |
|  |  |  |  | Average | stdev | Average | stdev | Average | stdev | Average | stdev | Average | stdev | Average | stdev |
| CZ | CV1 | CZ02, 06, 09, 10, 11 | 5 | 3,795,339 | 578 | 3,144 | 2.55 | 66.06 | 0.001 | 99.9987 | 0.0004 | 99.53 | 0.22 | 106.7 | 52.74 |
|  |  |  | (10) | 3,795,102 | 1,379 | 3,154 | 5.23 | 66.06 | 0.003 | 99.999 | 0.0005 | 99.50 | 0.03 | 96.6 | 57.93 |
|  | CV2 | CZ01, 14, 15, 18, 19, 20* | 6 | 3,791,236 | 11,241 | 3,169 | 57.03 | 66.06 | 0.013 | 99.9977 | 0.0021 | 99.24 | 1.17 | 230.07 | 226.58 |
|  |  |  | (10) | 3,795,535 | 1,309 | 3,158 | 4.12 | 66.06 | 0.002 | 99.999 | 0.0006 | 99.46 | 0.20 | 96.2 | 32.34 |
|  | CZ unique | CZ03, 04, 05, 13, 20, 22, 27, 29, 30, 31, 32, 34, 35, 36, 37 | 15 | 3,644,174 | 92,085 | 3,443 | 60.36 | 65.68 | 0.184 | 98.3352 | 0.2337 | 70.58 | 2.54 | 58,513.50 | 6,378.36 |
| CM | CM unique | CM02, 05, 06, 07, 08, 09, 10, 11, 12 | 9 | 3,823,282 | 69,250 | 3,350 | 60.31 | 66.01 | 0.123 | 98.5508 | 0.3637 | 70.82 | 2.62 | 54,261.1 | 9,340 |
| DZ | DZ unique | DZ01, 04, 05, 06, 07, 09, 10, 11, 12 | 9 | 3,630,009 | 91,461 | 3,425 | 147.4 | 66.01 | 0.253 | 98.2728 | 0.2383 | 71.64 | 2.22 | 58,115.9 | 7,089.9 |
| DW | CV3 | DW07, 11 | 2 | 3,710,853 | 623 | 3,208 | 3 | 65.81 | 0.017 | 99.9991 | nd | 99.59 | nd | 107 | nd |
|  |  |  | (4) | 3,710,023 | 348 | 3,213.5 | 6.24 | 65.81 | 0.001 | 99.9991 | 0.0005 | 99.11 | 0.62 | 82 | 32.52 |
|  | DW unique | DW07, 08, 10 | 3 | 3,688,086 | 17,366 | 3,142.7 | 45.44 | 65.91 | 0.168 | 99.9666 | 0.0108 | 94.23 | 2.18 | 2,924.33 | 1188.76 |
| UZ | CV4 | UZ13, 14 | 2 | 3,879,012 | 392 | 3,254 | 0.50 | 65.96 | 0.005 | 99.9994 | nd | 99.42 | nd | 82 | nd |
|  |  |  | (4) | 3,878,572 | 924 | 3,266 | 4.71 | 65.96 | 0.004 | 99.9991 | 0.0008 | 99.36 | 0.16 | 98 | 12.02 |
|  | UZ unique | UZ01, 02, 03, 04, 05, 06, 08, 09, 10, 11, 12, 13 | 12 | 3,771,146 | 141,577 | 3,642 | 169.52 | 65.47 | 0.230 | 98.2702 | 0.2615 | 69.61 | 4.33 | 60,896.40 | 8,666.32 |
| UM | UM<br>unique | UM08, 10, 12 | 3 | 3,765,537 | 89,752 | 3,335 | 120.33 | 66.01 | 0.292 | 98.26 | 0.5111 | 71.81 | 6.22 | 65,241 | 16,158.7 |
| FV | CV5 | FV6, 30, 23, 66, 69 | 5 | 3,732,326 | 13,119 | 3,181 | 20.02 | 66.12 | 0.019 | 99.9976 | 0.0007 | 99.63 | 0.07 | 88.1 | 22.66 |
|  |  |  | (10) | 3,728,750 | 10,824 | 3,185 | 10.99 | 66.12 | 0.019 | 99.999 | 0.0003 | 99.61 | 0.06 | 81.6 | 28.12 |
|  | CV6 | FV16, 35, 73, 68 | 4 | 3,811,847 | 1,478 | 3,250 | 2.21 | 65.97 | 0.002 | 99.9978 | 0.0006 | 99.57 | 0.07 | 103.67 | 51.12 |
|  |  |  | (8) | 3,811,276 | 1,621 | 3,260 | 4.37 | 65.96 | 0.003 | 99.998 | 0.0031 | 99.61 | 0.09 | 91 | 18.85 |
|  | FV unique | FV4, 5, 6, 8, 18, 41, 43, 53, 55, 60, 67, 70 | 12 | 3,802,109 | 59,614 | 3,303 | 122.72 | 65.88 | 0.168 | 98.3352 | 0.0021 | 70.58 | 2.54 | 62,206 | 16,720.50 |

\*The genome was not used for subsampling due to low coverage
nd the standard deviation was not calculated given that only two genomes were representing the CV

### Comparisons of genomic relatedness among the isolates reveals an intra-species ANI gap

The collection of 138 *Sal. ruber* genomes recovered from Mallorca and Fuerteventura solar salterns showed an average ANI value of 98.33% (SD = 0.37%) and a shared gene content of 73.38% (SD = 6.05%) (Fig. 1). ANI comparisons between isolates collected from the same sample showed similar results to the complete genome dataset (Fig. S2 and S3, for Mallorca ponds and Fuerteventura solar salterns, respectively) and the Fuerteventura genomes did not cluster separately from the Mallorca genomes in the ANI space (e.g., Fig. 1), revealing that similar *Sal. ruber* populations were present in the two distantly located islands (central-east Atlantic vs. northwest Mediterranean Sea; ~2000 km geographic distance). From the 9,454 ANI pairwise comparisons among all genomes in total, 1.79% showed ANI ≥ 99.8%, 0.35% between 99.6% and 99.8%, and 4.04% comparisons fell between 99.0% and 99.6%. The majority of ANI values (93.81%) fell between 97.0% and 99.0%. Therefore, the distribution of ANI values revealed an intriguing pattern, with low frequency of ANI values between 99.6% and 99.8% accumulating only 0.35% of the total comparisons vs. 1.24% expected if the total ANI values higher than 99% were uniformly distributed between 99% and 100%; that is ~4 times fewer values than expected by chance alone (Fig. 1).

**Figure 1:**
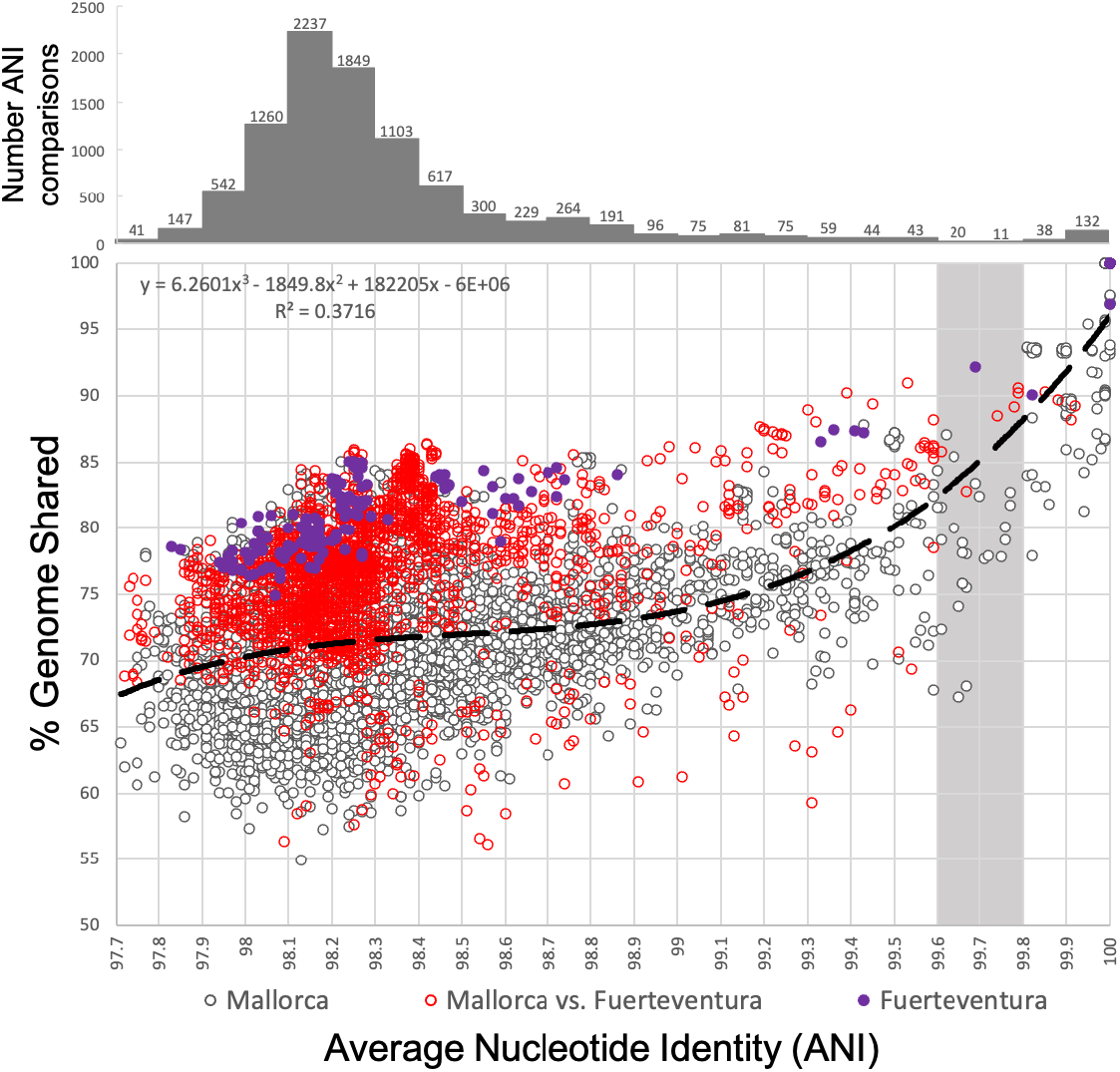
Genomic diversity of *Salinibacter ruber* genomes used in the study in terms of ANI relatedness and shared genome fraction. Each datapoint represents a comparison between two genomes and shows their ANI value (x-axis) against the shared genome fraction (y-axis). The graph on the top shows the number of datapoints for x-axis (in 0.1% windows or bins). Data points represent the 138×138 comparisons of our *Sal. ruber* isolate draft genome collection from Mallorca (118 genomes) and Fuerteventura (20 genomes) saline ponds combined (see also figure key for distinguishing datapoints by the place of isolation of the genomes compared). Note that the diversity of Fuerteventura genomes (in terms of ANI and shared gene content) is similar to that of many Mallorca genomes, albeit the latter collection also includes several more divergent genomes, in addition. Note also the shortage of data points (i.e., a gap) in ANI values around 99.6-99.8% (gray shaded area).

The shared gene content appears to follow ANI values, meaning that gene-content differences are typically larger among genomes with lower ANI values, but the relationship is biphasic. Genomes showing ANI >99.9% tend to have small gene-content differences (<5% of the total genes in the genome differ, on average). Differences in gene content increase sharply among genomes related between 99.5% and 99.9% before gene content differences stabilize around 15-35% for genomes showing <99.5% (Fig. 1). We examined the genomes that fell within the 99.6% and 99.8% ANI gap to gain a more quantitative view of the gene functions that differed at this level (Supplementary material). Our evaluation showed that the latter genomes had substantial gene content differences that could underlie ecological differences and/or adaptations such as phage predation, albeit typically smaller than those observed between more divergent genomes (showing ~98% ANI) and larger than those observed between closely related genomes, ANI >99.8%.

### A natural definition for a genomovar and strain

The ANI gap revealed around 99.6-99.8% ANI (Fig. 1) was the only ANI range with such a strong bias in terms of pairwise comparisons not falling in this range [Gap here refers to the small number of genome pairs showing 99.6-99.8% ANI relative to counts of pairs showing ANI >99.8% or <99.6%]. It is unlikely that this ANI gap is due to cultivation biases because the media and conditions used are thought to be robust for *Sal. ruber* and do not distinguish between members of the species or closely related *Salinibacter* species (19, 22). Therefore, the 99.6-99.8% ANI gap appears to be a property emerging from the data itself and to be robust. We suggest that the term genomovar could be used to refer to these ANI-based intra-species units. The term genomovar was originally used to name distinct genomic groups within species that cannot be phenotypically differentiated and therefore, cannot be classified as distinct species based on the standard (low-resolution) taxonomic practices of the past (25). The original definition was tailored “*to allow a nomenspecies to contain more than one genomic group, … analogous to other subdivisions of the nomenspecies”* (25). The suffix-var indicates an intra-specific subdivision (10), not covered by the bacteriological code, but recommended to be used to describe intra-species groups. Hence, genomovar may capture well the intra-species ANI gap revealed by our analysis. We also propose the lower value of the ANI gap revealed (~99.5% ANI) as opposed to the upper value (99.8% ANI) of the gap as a more conservative threshold to define genomovars. For the strain level, 0.5% or 0.2% difference in ANI (correspondingly, 99.5% and 99.8% ANI) represents substantial, non-trivial, genomic divergence that, in most cases, would likely encompass several genomes with at least some phenotypic differences (due to substantial sequence or gene content differences among the genomes; see Fig. 1). Thus, multiple strains will be likely grouped together under the same 99.5% ANI cluster in such cases, and strain, in general, represents a more fine-grained level of resolution than the 99.5% ANI level.

Accordingly, we propose to define strain as a group of isolates showing ANI values >99.99%. This threshold also ensures high gene content similarly; e.g., shared genome usually >99.0% based on the *Sal. ruber* genomes analyzed that show >99.99% ANI, and encompasses well the typical sequencing and assembly noise observed. For instance, to quantify the noise resulting from the high-draft status (incomplete) of our genome sequences, we selected 23 isolates with sequencing depth >100X, split the raw data in two halves, and assembled the two subsets independently for direct pairwise comparisons of the resulting genome sequences. The ANI value between the paired re-assembled genomes showed values >99.99% and shared genome fraction >99.1% (Table 1; Sup. Spreadsheet S1). Finally, a strain should not be equated to clone as the latter implies identical sequence, and the definition of strain may allow a certain degree of genome and gene-content divergence (25). Consistent with this assumption, all but two genomes that were assigned to the same CV based on identical RAPD profiles also showed ANI values >99.99%. Thus, our proposed definition for strain is compatible with CVs as the latter have been traditionally defined (discussed further below).

### Relative abundance of isolates in the samples of origin reveals cultivation biases

Next, we assessed what fraction of the total *Sal. ruber* population in the sample of origin each isolate made up based on the number of metagenomic reads competitively recruited by the isolate-specific genes as well as (independently) the core (shared) genes to assess the magnitude of isolation biases. Given the high ANI identities among any isolate genome in our collection (>97%) and the distribution of nucleotide identities of shared genes among the isolates (see also below), we defined the *Sal. ruber* population as any metagenomic sequence sharing >95% nucleotide identity with any genome sequence in our collection. To assess the feasibility of our core gene-based approach, we first examined the nucleotide identity patterns among alleles of core-orthologous genes within vs. between genomovar comparisons. From the pangenome analyses, including all sequenced genomes from Mallorca and Fuerteventura (138 isolates), we detected 793 core single-copy orthologous groups (core-OGs: genes encoded in all genomes without paralogs). We considered an allele of a core gene to be unique when showing identity <99.8% to other alleles of the same gene based on the intra-vs. inter-genomovar ANI values, which also allowed for 1 to 2 sequencing and/or assembly errors resulting in SNVs that are common at the edges of assembled contigs. Our analysis revealed a total of 26,325 unique alleles among the 793 genes shared by our isolate genomes. About 66% of the core-OGs (523 out of 793) showed between 22 and 37 different alleles each (Fig. S4), and we identified two low-diversity core-OGs (high sequence conservation) that encoded for only ten alleles (ribosomal protein L32 and an ATP-binding cassette protein) and two highly diverse core-OGs with 111 different alleles each (cold shock protein and conserved hypothetical proteins). In general, all isolates belonging to the same genomovar shared always 100% of the core-OG alleles (Fig. S6) and an average of only 9.7% (SD = 14.5%) of the alleles were shared between different genomovars (Sup. Spreadsheet S2). Further, and as expected, the clustering based on the percentage of shared alleles between genomes and the phylogeny based on the concatenated sequence alignment of the 793 core-OGs were in good agreement (Fig. S5). Therefore, it appears that allelic variation in OGs between genomovars was generally adequate to distinguish genomovars (but not isolates or strains within genomovars), especially if metagenomic reads are recruited to representative genomes of genomovars (one representative per genomovar) competitively, albeit with some saturation in the signal (e.g., identical OG shared between genomovars). We dealt with ties in identity in read mapping by not counting such reads.

Interestingly, analysis of the results from competitive recruitment of *Sal. ruber* reads against representative genomes of genomovars showed that the most commonly recovered genomovars based on the number of isolates assigned to them were not the most abundant genomovars in their natural populations (Table S1). For instance, from the 16 different isolates recovered at time zero from the control pond of Mallorca, the most abundant genomovar-CV1 and genomovar-CV2 (represented by 5 and 6 isolates, respectively; genomovars are named after the most abundant, or only CV, its isolates were assigned to) were the least abundant in the metagenome (0.37% of the total the *Sal. ruber* population), and the isolates CZ13 and CZ31, each assigned to a different single-isolate CV, represented the most abundant genomovars *in-situ* with 2.86% and 1.74% relative abundance, respectively. Similarly, from another pond [short-shaded pond at the initial time of the experiment, which is similar to the control pond because the shade treatment had not been applied to it yet (23)] two isolates were assigned to genomovar-CV4, which showed a relative abundance of 0.98%, while the most abundant genomovars *in-situ* were the (single-isolate) UZ12, UZ02, and UZ08, which showed a relative abundance of 3.03%, 2.76% and 2.62%, respectively. The results were similar between mapping to core vs. all genes in the genome (Fig 2). The same analysis applied to the Fuerteventura metagenome revealed that the nine isolates represented by genomovar-CV5 and genomovar-CV6 (relative abundance of 10.6%) showed lower abundance than the (single-isolate) FV41 and FV43 (18.3% and 28.3% respectively).

**Figure 2:**
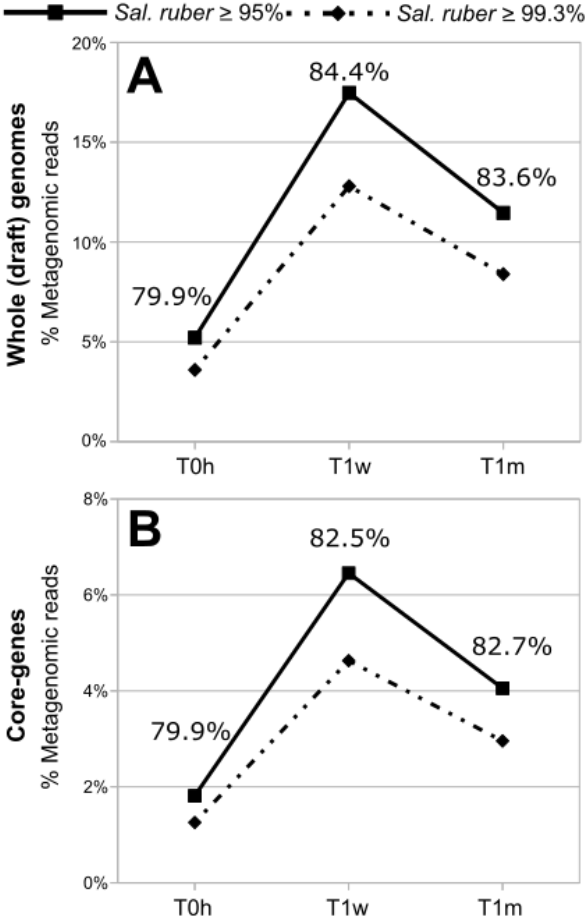
*Salinibacter ruber* abundance dynamics in the control pond during the sampling period of one month. Metagenomic reads were aligned against *Sal. ruber* genomes using (A) whole (draft) genomes and (B) core genes only (see text for details). Continuous lines show the abundance of the natural *Sal. ruber* population (represented by reads mapping with identity ≥95% to any genome) and discontinuous lines show the percentage of reads represented by the sequenced isolates (reads mapping with identity ≥99.3% to any genome). Numbers for each time point indicate the proportion of total population reads that were associated to sequenced strains.

### Estimations of the total number of genomovars making up the natural *Sal. ruber* population

The *Sal. ruber* population showed a relative abundance ranging between 5.7% and 19.5% of the total microbial community represented in the control (no treatment) pond metagenomes of the Mallorca salterns, depending on the sample considered, and 6.12% of the total microbial community in the sequenced Fuerteventura metagenome sample. To estimate the level of coverage of the intra-population sequence and gene-content diversity achieved by our sequencing effort, we applied the read redundancy approach of Nonpareil (26) to the metagenomic reads identified as *Sal. ruber* (≥95% identity against any of the genomes in our collection) with a pairwise read identity threshold of ≥99.3% to identify redundant reads. The approach posits that if the reads are completely redundant, then the sequencing has saturated the extant intra-population diversity. We used 99.3% in this case to allow for one mismatch or sequencing error given our average read length of 150 bp, e.g., 149/150=99.33. Using 100% identity did not differentiate our conclusions substantially (data not shown). Note also that the 99.3% identity threshold is close to the 99.5% ANI threshold that distinguishes genomovars; thus, its use should not confound results from competitive read mapping against isolates of different genomovars. The estimated coverage achieved in the Mallorca control metagenomes when the three control samples were combined ranged between 93% and 95% (Table S2). These results indicated that our reads, collectively, sampled the great majority of the intra-population diversity of the *Sal. ruber* present in the samples, which was also consistent with the high relative abundance of the population *in-situ* reported above. Nonpareil’s sequence diversity *Nd* index also indicated that the intra-population diversity decreased only slightly during the sampling period and by the treatments, from 14.58 at time zero to 14.47 at time one month (*Nd* is in natural log scale; Table S2).

Metagenomic reads mapping at high identity (≥99.3%) to an individual *Sal. ruber* isolate genome representing a distinct genomovar captured between 30% and 40% of the *Sal. ruber-identified* reads from Mallorca, depending on the sample and the isolate considered, and reflecting both genomovar-derived sequences as well as highly conserved genes across genomovars such as those originating from the rRNA gene operon (noise). Comparing the percentage of metagenomic reads assigned to the *Sal. ruber* species (identity ≥95%) and those mapping with high identity to genomovars represented by isolates in our collection (identity ≥99.3%), our collection of sequenced isolates recruited 79.9% of the species reads at time zero, 84.4% at one week, and 83.6% at one month of the corresponding samples from the Mallorca control pond (salt saturation conditions). Similarly, the 12 sequenced genomovars (20 isolate genomes, in total) represented a total of 77.3% of the metagenomic reads assigned to *Sal. ruber* for the Fuerteventura metagenome (Fig. 3). Therefore, about 20% of the reads assigned to *Sal. ruber* were not recruited by our isolate collection, indicating that a higher effort in cultivation is necessary to fully recover all abundant genomovars of the population.

**Figure 3:**
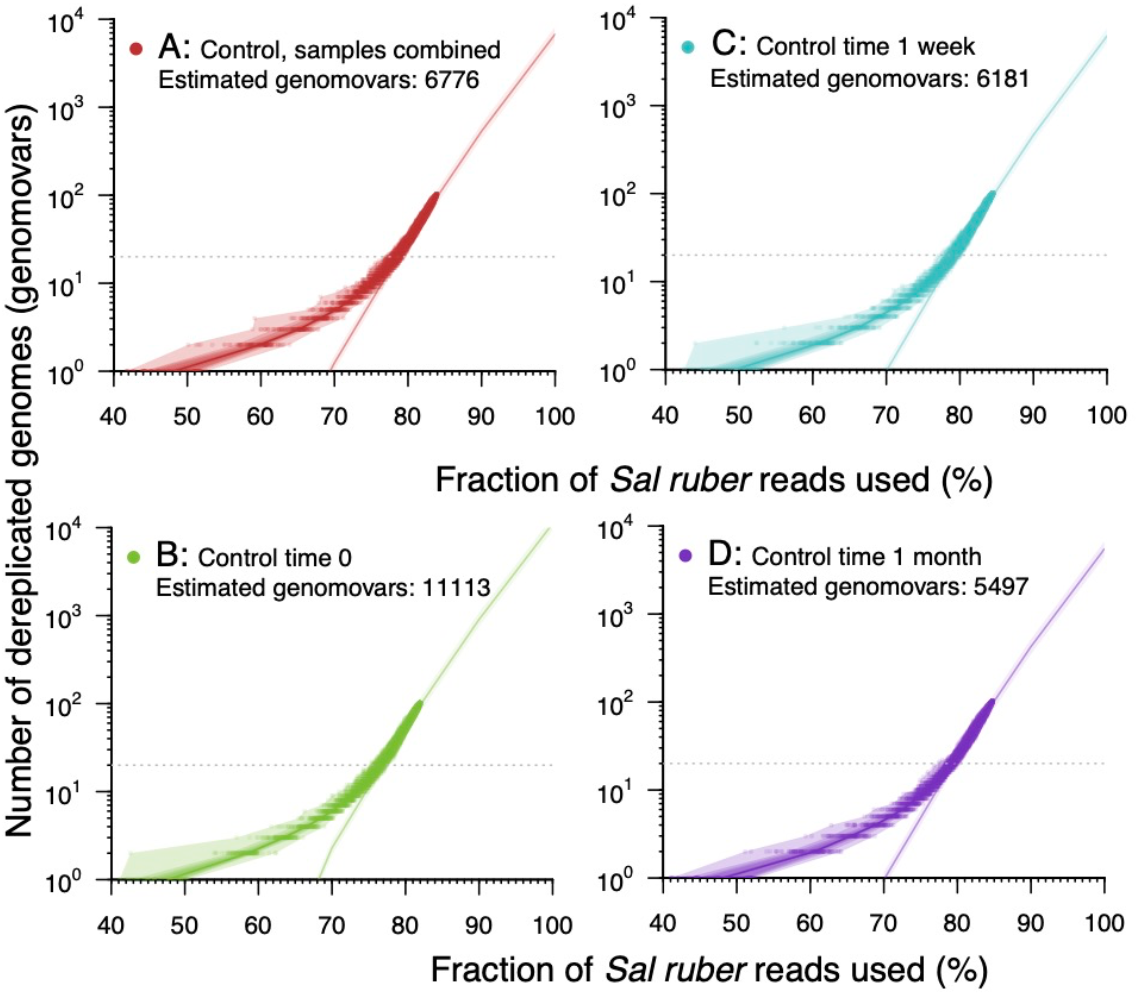
Estimation of the number of genomovars making up the natural *Salinibacter ruber* population. Metagenomic reads from each sample (Panels B-D) or all samples combined (Panel A) of the control pond were mapped to the *Sal. ruber* genomes preserving all matches with identity ≥99.3%. The mapping file was manipulated to remove one target genome at a time (randomly sorted) while recording the number of unique reads mapping at each step, and this process was repeated 100 times to reduce the impact of randomization on the estimates obtained (below). The number of reads were then expressed as the fraction of the maximum number of reads from the *Sal. ruber* species by dividing the observed counts by the total number of reads mapping to any reference genome with identity ≥95%. The logarithm of the number of total (dereplicated) genomes used was then expressed as a function of the fraction of *Sal. ruber* reads captured by the genomes, and a linear regression was determined by unweighted least squares and evaluated using Pearson correlation for the region between 20 and 100 genomes. This trendline was extrapolated to 100% coverage of the genomovar diversity (i.e., all reads from the species) to provide an estimate of the number of genomovars represented (Y-axis) in the total sequenced fraction (X-axis). Filled dots represent the fraction of the total *Sal. ruber* reads captured by the genomovars used, and the shaded bands around the observed subsamples represent the central inter-quantile ranges at 100%, 80%, 60, 40%, and 20%.

To evaluate the number of genomes that will be necessary to sequence in order to recover the full *in situ* species genome diversity (at the genomovar level, since we cannot reliably resolve between genomes or strains of the same genomovar) using the same isolation procedure as applied herein, we built a rarefaction curve of the number of reads recruited by each isolate against the number of isolates included in the analysis (one isolate genome per genomovar was used in the analysis). We observed that, using a logarithmic scale (Fig. 3) between 20 and 100 genomes, the rarefaction curve is almost a straight line (Pearson correlation value of 0.99935), and we would need to isolate and sequence around 11,000 isolates (99% prediction interval: 9,205 – 13,416), each representing a distinct genomovar, in the more diverse sample (control time zero) and around 5,500 isolates (4,501 – 6,711) in the lower diversity sample (control time 1 month) in order to capture the complete extant genetic diversity (Fig. 3 and Table S3).

### RAPD genotyping is a reliable method for identifying redundant genomes or members of the same strain

The available genomes allowed us to assess the resolution of the RAPD method for identifying clonal isolates based on direct comparisons to the genome sequences of the corresponding isolates. In general, we observed that almost all isolates sharing identical RAPD patterns using three different PCR primers also showed ANI values >99.99% and shared genome >99.24% while genomes of different RAPD patterns (different CVs) typically showed ANI values <99.8% and shared genome <96%. Accordingly, the genome size between isolates of the same CV varied only slightly, between 392 bps in CV4 and 13,119 bps for CV5 (Table 1; Sup. Spreadsheet S3). We identified only two exceptions to these patterns. Specifically, although the nine sequenced isolates from Fuerteventura solar salterns displayed an identical RAPD pattern, we decided to split the genomes in two different CVs (CV5 and CV6) based on genomic differences (e.g., ANI were <99.90% and 70 genes consistently differed between isolates of CV5 vs. CV6). Further, although the isolates assigned to CV1 and CV2 showed different RAPD patterns based on RAPD4 primers, their genomes displayed very high similarity, comparable to that observed in genomes within other CVs (Sup. Spreadsheet S4). The ANI between members of CV1 and CV2 was 99.99% on average (SD = 0.0012%) and percentage of shared genomic fraction 99.38%, on average (SD = 0.41%). We were not able to detect any substantial difference based on the genome comparison and thus, merged the two CVs into one CV. In summary and based on our proposed definition for strain (>99.99% ANI and >99.0 shared gene-content) and analysis of 138 isolates genomes, in >95% of the cases the RAPD methodology was accurate in assigning a genome to the same (or different) strain based on identical (or non-identical) profiles. Thus, RAPD profiling represents a quick, cost-effective and reliable method for genotyping isolates based on genomic evaluation, which is consistent with previous results based on low-resolution fingerprinting methods (24).

## DISCUSSION

Distinct units within bacterial species have been recognized long time ago, and have been designated with different names such as subspecies, ecotypes, clonal complexes, serotypes, genomovars and strains, among several other designations [reviewed in (27)]. However, the use of these units has commonly been inconsistent between different studies, and the units are often not applied based on the same standards or means (e.g., marker genes used) across different bacterial taxa, creating challenges in communication about diversity. In a comparison of 138 *Sal ruber* isolates we observed an area of genomic discontinuity between 99.6% and 99.8% ANI that accumulated only about one quarter of the total pairwise measures expected by chance in case of a uniform ANI value distribution. This natural gap emerging from the data itself, as opposed to a manmade arbitrary threshold, led us to suggest the concept of genomovar (25) for all organisms sharing genome-wide ANI > 99.5%. Our proposal provides a more accurate and standardized definition of this unit compared to previous practice. Importantly, a recent analysis of 330 bacterial species with at least 10 complete (not draft) sequenced isolates per species available in the NCBI database showed that the 99.2-99.8% ANI gap is also found within most of these well-sampled species (28). Therefore, the genomic discontinuity around 99.5% ANI observed here based on the *Sal. ruber* genomes may be a universal property of bacterial species and thus, a robust, natural definition of genomovars within species.

Our analysis also revealed that there is significant ANI and gene content diversity within a genomovar, defined at the 99.5% ANI level. To circumscribe this diversity, we also propose ANI >99.99% as a general purpose and practical threshold to define a strain within a genomovar, which ensures high gene content similarity among members of the same strain (typically, >99.0% of total genes) and is robust to the noise of the genome sequencing and assembly processes (Table 1). Further, this threshold almost always corresponded to identical RAPD fingerprints, with only a few exceptions of genomes sharing >99.99% ANI that differed in their RAPD profile and/or the gene content substantially. For example, isolates of the CV6 RAPD group could be differentiated from CV5 isolates by just the presence of a plasmid in the former, which encoded for a CRISPR-Cas system thus, could provide an adaptatively immune function against viruses (29, 30). We propose to consider such cases of genomes having substantial gene-content differences attributed to mobile elements while sharing >99.99% ANI still as representative of the same strain; in other words, to let the ANI > 99.99% threshold override mobile element differences in order to simplify strain identification and communication. Further, mobile elements are often ephemeral and do not confer substantial phenotypic differences in cases that they are not expressed or carry functionally important genes. However, in cases where important phenotypic differences that distinguish between organisms sharing ANI >99.99% are known such as antibiotic resistance genes carried by plasmids, our proposed definition for strain could be neglected or adjusted to even higher ANI values as seen appropriate. It follows that a strain differs from the concept of a clone that implies identical genome sequence (Fig. 4).

**Figure 4:**
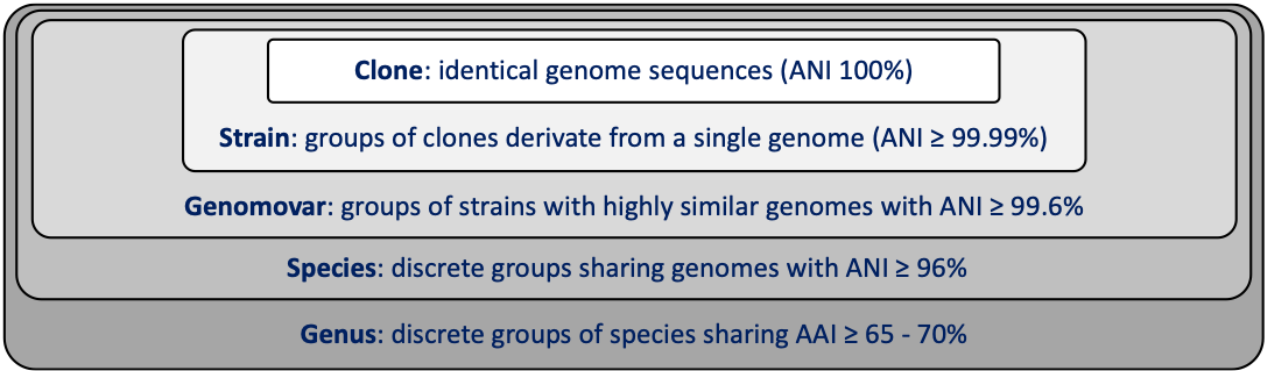
Proposed thresholds to define species and subspecies units. Note that thresholds are based on average amino-acid identity (AAI) for genus level and average nucleotide identity (ANI) for species and intra-species level.

Based on the abovementioned definitions and thresholds for genomovar and strain, our findings suggest that the number of (distinct) genomovars making up the *Sal. ruber* population appears to be large but not unfathomable. Specifically, our mathematical extrapolations indicated that to completely cover the genomic diversity of the natural population of *Sal. ruber*, we would need to isolate and sequence between 5,500 and 11,100 isolates, each representing a distinct genomovar; a large but bounded number. These results contrast with a previous estimation indicating that diversity within species may be unbounded (31). It should be mentioned, however, that since our sequencing effort did not saturate the diversity within the natural *Sal. ruber* population (~95% of diversity was covered by sequencing), it is possible that our estimates on the number of isolates needed might change with higher coverage, especially if the diversity not captured by our sequencing effort harbors a disproportionately lower (or higher) number of genomovars than are accounted by our fitted trendlines (e.g., Fig. 3). Furthermore, performing this type of analysis with long-reads, or even single-cell amplified genomes, could reduce uncertainty in the estimations caused by the use of short reads, which cannot resolve well very closely related genomes due to the limited sequence available. Most importantly, such long-read data will reveal whether or not the associated sequence thresholds for identifying redundant reads by our study underestimated the diversity of genomovars within the *Sal. ruber* population.

In contrast to the high genomovar diversity estimated, we find it remarkable that our relatively small collection of sequenced isolates (107 out of 187 unique RAPD profiles for the Mallorca ponds) already represented ~80% of the *Sal. ruber* population in the habitat, on average (Fig. 2). These results suggested that most genomovars not represented by our isolates had low abundance *in-situ*, at least at the time of our sampling. Therefore, it appears that the concept of the rare biosphere for the species or population levels (32) applies equally well to the genomovar abundance patterns within the natural *Sal. ruber* population. These results also suggested that our isolation procedures were overall robust and did not dramatically bias the diversity of the *Sal. ruber* population recovered, meaning that the genomovars that appear to be the most abundant *in-situ* were represented among the recovered isolates. Given that representative isolates of the abundant genomovars were recovered from the control as well as the treatment pond samples for sunlight reduction (shade application) and brine salt dilution, the two major environmental drivers of the saltern systems, our results further indicate that the genomovars represent mostly neutral diversity. That is, most of them should be functionally redundant; a hypothesis that remains to be more rigorously tested in the future.

Despite the overall high representation of the *Sal. ruber* extant diversity by our isolates, however, the most frequently recovered genomovars from a single sample were frequently not the most abundant genomovars *in-situ*. In fact, it appeared that the most abundant genomovars by isolation often were among the least abundant *in-situ* genomovars among those represented by our isolate collection based on competitive read mapping to isolate/genomovar-specific or core genes. This result might not be surprising in the *Sal. ruber* case that most genomovars or strains recovered appear to be relatively low-abundance *in-situ* according to our analysis (Fig. 3). Further, cultivation biases are well appreciated in microbial ecology for several decades now (33). Nonetheless, our results do show that the laboratory growth conditions could select for less abundant members of a population (i.e., lead to cultivation biases), even in cases where the growth media may be specifically-designed for the target population and appear to be robust, in general. Therefore, for reliable estimates of *in-situ* abundance, isolation efforts need to be combined with culture-independent data even in cases where growth media are thought to not be restrictive for the target organism. Recovery of a higher number of isolates could also help with cultivation biases, e.g., despite our substantial efforts only a small number of isolates were members of the same genomovar due to the high intra-species diversity, albeit obtaining a higher number of isolates than what was achieved here (n=409) may be impractical and/or costly.

The thresholds proposed here (Fig. 4) to define genomovars and strains provide convenient and practical means to define these sub-species units and thus, should greatly facilitate future studies in environmental or clinical settings. We also suggest evaluating the ANI value distribution for the species of interest, and if the data indicate so, to adjust the ANI threshold to match the gap in the observed ANI value distribution. Finally, our results on the diversity and number of genomovars found within a natural population as well as the extent of cultivation biases should represent a useful guide for future microdiversity surveys.

## Materials and Methods

The Supplementary Material includes additional details to those provided in the Results section about how genome and metagenome data were obtained and bioinformatically processed as well as details about the samples obtained and how isolates were typed based on RAPD and MALDI-TOF MS.

## Data availability

Genomes and metagenomes sequenced for this study from Mallorca and Fuerteventura solar salterns have been deposited in the European Nucleotide Archive (ENA) under BioProject accession numbers PRJEB27680 and PRJEB45291, respectively (Sup. Spreadsheet S5).

## Acknowledgements

The authors would especially like to thank the whole team at Salinas d’Es Trenc and Gusto Mundial Balearides, S.L. (Flor de Sal d’Es Trenc) and of the Salinas de Fuerteventura (Salinas del Carmen) for allowing access to their facilities and their support in performing the experiments. This study was funded by the Spanish Ministry of Science, Innovation and Universities projects MICROMATES PGC2018-096956-B-C41 and PGC2018-096956-B-C43, MARBIOM RTC-2017-6405-1 and METACIRCLE PID2021-126114NB-C42 and PID2021-126114NB-C44, which were also supported by the European Regional Development Fund (FEDER). KTK’s research was supported, in part, by the U.S. National Science Foundation (Award No. 1831582 and No. 2129823). RRM acknowledges the financial support of the sabbatical stay at Georgia Tech supported by the grant PRX18/00048 of the Ministry of Sciences, Innovation and Universities. The computational results presented here have been achieved (in part) using the LEO HPC Infrastructure of the University of Innsbruck. The Fuerteventura samples were taken in accordance with the permit ESNC27, with the unique identifier ABSCH-IRCC-ES-241224-1 that has been provided by the Dirección General de Biodiversidad y Calidad Ambiental del Ministerio para la Transición Ecológica of the Spanish Government. TV acknowledges the “Margarita Salas’ postdoctoral grant, funded by the Spanish Ministry of Universities, within the framework of Recovery, Transformation and Resilience Plan, and funded by the European Union (NextGenerationEU), with the participation of the University of Balearic Islands (UIB). TV and RA acknowledge support by the Max Planck Society.

