## Supplementary material for "Towards estimating the number of strains that make up a natural bacterial population"

**Running Title:** How many strains make up a bacterial population?

**Affiliations**

[1] Marine Microbiology Group, Department of Animal and Microbial Biodiversity, Mediterranean Institute for Advanced Studies (IMEDEA, CSIC-UIB), Esporles, Spain.

[2] Department of Molecular Ecology, Max Planck Institute for Marine Microbiology, Bremen, Germany.

[3] School of Civil and Environmental Engineering, and School of Biological Sciences, Georgia Institute of Technology, Atlanta, GA, USA.

[4] Department of Microbiology, and Digital Science Center (DiSC), Universität Innsbruck, Innsbruck, Tyrol, Austria.

[5] Unidad de Epidemiología y Medicina Preventiva, IUSA, Facultad de Veterinaria, Universidad de Las Palmas de Gran Canaria, C/Trasmontaña s/n, Arucas, 35413, Canary Islands, Spain.

[6] Department of Biochemistry, Genetics and Microbiology, and Forestry and Agricultural Biotechnology Institute (FABI), University of Pretoria, Pretoria, South Africa.

[7] Department of Mathematics and Computer Science, University of the Balearic Islands, Palma 07122, Spain.

**Materials and methods:**

***Experimental description***

The *Sal. ruber* isolates analyzed here were collected from six adjacent solar saltern ponds located in Mallorca (Spain) that have been previously used for manipulative experiments described in (1, 2). In addition, as part of the present study, we included one metagenome sample from Fuerteventura saltern (Salinas del Carmen) located in Canary Islands (Spain) collected in 2020 and 40 isolates that originated from this single sample. The Fuerteventura sample was collected in accordance with the permit ESNC27, with the unique identifier ABSCH-IRCC-ES-241224-1 that has been provided by the Dirección General de Biodiversidad y Calidad Ambiental del Ministerio para la Transición Ecológica of the Spanish Government.

Isolates from these samples were recovered in sea water medium supplemented with 25% salt concentration (3) and taxonomically identified by Matrix-Assisted Laser Desorption Ionization-Time of Flight Mass Spectrometry (MALDI-TOF MS) as described in (4). RAPD fingerprinting (5) was applied to identify clonal isolates. RAPD fingerprints were obtained using the RAPD4, RAPD5, and RAPD6 amplification primers (6).

***Genomic sequencing and analysis***

DNA extraction was performed as detailed in (7). DNA sequencing libraries were prepared using the Illumina Nextera XT library prep kit and libraries were sequenced using a NovaSeq6000 150PE (2 x 150bp) instrument. Raw reads were trimmed, and quality filtered to remove low quality sequences using BBDuk v38.82 (http://bbtools.jgi.doe.gov). Options used for trimming were: ktrim=r, k=28, mink=12, hdist=1, tbo=t, tpe=t, qtrim=rl, trimq=20 and minlength=100. Trimmed reads were assembled using SPADES v3.14 (8) with default parameters. Gene prediction using contigs longer than 500 bp was conducted using Prodigal v2.6.3 with default parameters (9). Genes were annotated using SwissProt and TrEMBL databases (10) with Diamond tool using default settings (11). Results were filtered for best match based on bit score, using a minimum threshold of 40% sequence identity and 50% match length. The ANI value between all vs. all genomes was determined using the *ani.rb* script from the

Enveomics collection (available at http://enve-omics.gatech.edu) (12). SNVs detection on assembled contigs was performed using Mauve v2.4.0 (13).

Pangenome analysis was applied using gene sequences at the nucleotide level. Predicted gene sequences were compared using an all-versus-all BLASTn v2.2.28 (14) to identify the shared reciprocal best matches based on bit score in all pair-wise genome comparisons using a 90% sequence identity cut-off and 90% or more coverage of the query sequence length. The identification of Orthology Groups (OGs), defined as the reciprocal best matches among genomes, was performed using the *ogs.mcl.rb* script from the Enveomics collection (12). Our core-genome analysis identified 797 shared (core) OGs in single copy (no apparent paralogues). These core-genes were aligned individually using muscle aligner v3.8.31 in order to build consensus trees (15). For the latter, aligned genes were concatenated using the *Aln.cat.rb* script from the Enveomics collection (12) and phylogenetic analysis was performed using the Neighbor Joining algorithm implemented in ARB v6.0.6 (16). The collection of universal/housekeeping genes were extracted using the *HMM.essential.rb* script from the Enveomics collection (12).

***Metagenome sequencing and analysis***

Metagenomes from Mallorca salterns were obtained as detailed in (17). DNA extraction of the sample collected from the Fuerteventura saltern was performed as detailed in (7). DNA sequencing libraries were prepared and sequenced as described above for isolate genome libraries. Metagenomics reads were quality-trimmed as described above and subsequently, mapped against all *Sal. ruber* genome sequences competitively (meaning, only the best matching genome among all possible genome matches for each read was retained, and only if above the cut-off for a match; ties for best matches were not counted), using BLASTn v2.2.28 (14), to identify reads belonging to each genome or its very close (not sequenced) relatives. After the Blast search, best match read selection (based on bit score) was determined using the *BlastTab.best_hit_sorted.pl* script from the Enveomics script collection (12).

To calculate the total abundance and intra-population diversity of the *Sal. ruber* natural population, we selected reads mapping with sequence identity ≥95% and read coverage ≥70% to any *Sal. ruber* genome in our collection. Subsets of these reads mapping with identity ≥99.3%

(using BLASTn (14) as described above; note that 1 mismatch in reads of 150 bps in length results in 99.3% identity) were selected to calculate the abundance of individual isolates or genomovars. The total number of reads mapping to any *Sal. ruber* genome (at identity ≥95%) were normalized by the total sequencing effort (i.e., total number of reads per metagenome) to obtain the final relative abundance for the *Sal. ruber* population (or species). To calculate the difference in abundance between different isolates or genomovars, the analysis only considered the reads that mapped at identity ≥99.3% to one genome and <99.3% to the remaining genomes when representing different genomovars compared to the former genome. The number of reads mapping to each genome was divided by both genome length and total number of metagenomic reads to provide the final relative abundance for the corresponding genome, and by extension, the genomovar the genome represented.

To calculate the coverage of the total *Sal. ruber* population obtained by the sequencing effort applied, we used the Nonpareil tool v3.303 (18) on metagenomic reads mapping with identity ≥95% and ≥70% coverage with any of the genomes in our collection. Options used for Nonpareil analysis were: Nonpareil algorithm (-T) = alignment, maximum number of reads as query (-X) = 1,000,000 and identity threshold to group two reads together (-S) = 0.993.

To calculate the rarefaction curves of the genomovar diversity captured by the sequencing effort applied, metagenomic reads were mapped to *Sal. ruber* genomes preserving all matches with identity ≥99.3%. The mapping file was manipulated to remove one target genome at a time (randomly sorted) while recording the number of unique reads mapping at each step, and this process was repeated 100 times to reduce the impact of randomization on the estimates obtained (below). The number of reads were then expressed as the fraction of the maximum number of reads from the *Sal. ruber* species by dividing the observed counts by the total number of reads mapping to any reference genome with identity ≥ 95%. The logarithm of the number of total (dereplicated) genomes used was then expressed as a function of the fraction of *Sal. ruber* reads captured by the genomes, and a linear regression was determined by unweighted least squares and evaluated using Pearson correlation for the region between 20 and 100 genomes. This trendline was extrapolated to 100% coverage of the genomovar diversity (i.e., all reads from the species) to provide an estimate of the number of genomovars represented in the total sequenced fraction. The linear regression, together with the prediction interval, were calculated with the R package stats v4.0.3. The results are reported in the main text, Figure 3, and Suppl. Table S3.

**Supplementary Text S1: Genomic differences between isolates of distinct CVs.**

All members of the same CV consistently exhibited ANI >99.99%, but we also observed a few cases of genomes belonging to different RAPD categories that showed high ANI values. For example, CV3 isolates (DW07 and DW11), despite being distinct based on RAPD4, showed highly similar RAPD6 profiles with isolates DW08 and DW10 obtained from the same sample. Moreover, RAPD5 discriminated between CV3 (DW07 and DW11) and both DW08 and DW10, which shared identical pattern (Fig. S7). Consistently, genomes from CV3 displayed ANI values >99.96% with genomes DW08 and DW10, and an average shared genome fraction of 91% with DW08 and 93% with DW10. Genomes DW08 and DW10 displayed an ANI value of 99.94% and shared genome fraction of 95.4% between them (Sup. Spreadsheet S6). The pangenome of the four isolates indicated that the genomes from CV3 encoded for an identical set of 3,208 genes of which 258 were specific to CV3. 131 genes were specific to DW08 and 64 genes to DW10. Most strain-specific genes were encoded in a single contig in each respective assembly, indicating that they represent a genome island (e.g., a plasmid or a prophage). The functional annotation of CV3 genomes showed that among the 258 CV-specific genes there were a phage integrase, an intron reverse transcriptase, a DNA helicase, two different CRISPR Cas systems (one of them associated to CRISPR type I-E), a plasmid partition protein (ParA) and an environmental halophage (Sup. Spreadsheet S7). In genome DW10, the genome-specific genes encoded for a ParA, 5 transposases, 1 integrase, 1 helicase, and a pilin glycosylation enzyme, in addition to hypothetical and other poorly characterized proteins. In genome DW08, the genome-specific genes encoded for viral genes (as VirE and nucleases), phage genes Resolvase/invertase-type recombinase genes and three different contigs encoded for ParA genes.

Similarly, CV5 and CV6 isolates showed nearly identical RAPD profiles using the three set of primers. However, isolates from CV6 presented a genome length ~80,000 bps larger than those of CV5. Pan-genome analysis comparing these two CVs revealed that over 99.5% of the CV5 genes were shared with CV6 isolates. Specifically, CV6 carried 56 additional genes not carried by CV5. Analyzing the metabolic functions in the FV16 genome (CV6), we detected that these genes were carried by two contigs (contig 22 and 28; Sup. Spreadsheet S8); whereas the corresponding genes were in the same contig in genome FV68. In the two contigs, we detected the presence of CRISPR spacers. Contig 22 encoded for 2 nucleases, a DNA polymerase, 2 transposases, a DNA invertase, a recombinase and an endonuclease. Contig 28 encoded one ParA family protein, a RepB plasmid replication protein, and one CRISPR Cas system (Cas2, Cas1, Cas4, Cas7, Cas3, Cas6 proteins). The gene annotation indicated that the contigs belong to a plasmid and the coverage was two times higher than the chromosome indicating a double-copy plasmid. Altogether, these results showed that the most closely related CVs had substantial gene content differences (in addition to their ANI dissimilarities), albeit typically smaller than those observed between most CVs sharing ~98% ANI, that could underlie ecological differences and/or adaptations such as phage predation.

**Supplementary Figure S1: Identification of *Sal. ruber* clonal varieties (CVs) using RAPD signatures.** Identical RAPD profiles were interpreted to belong to the same CV. RAPD fingerprints were obtained using RAPD4, RAPD5 and RAPD6 amplification primers (6). Each column in the gels shows a RAPD profile for a different isolate. Isolates of CV1 to CV4 were retrieved from a mesocosm experiment from the Mallorca solar salterns (CZ: Control pond time zero; DW: Dilution pond time 1 week; UZ: Unshaded pond time zero). The mesocosm experiments were described previously (1, 2, 17). Isolates of CV5 and CV6 were retrieved from Fuerteventura solar salterns.

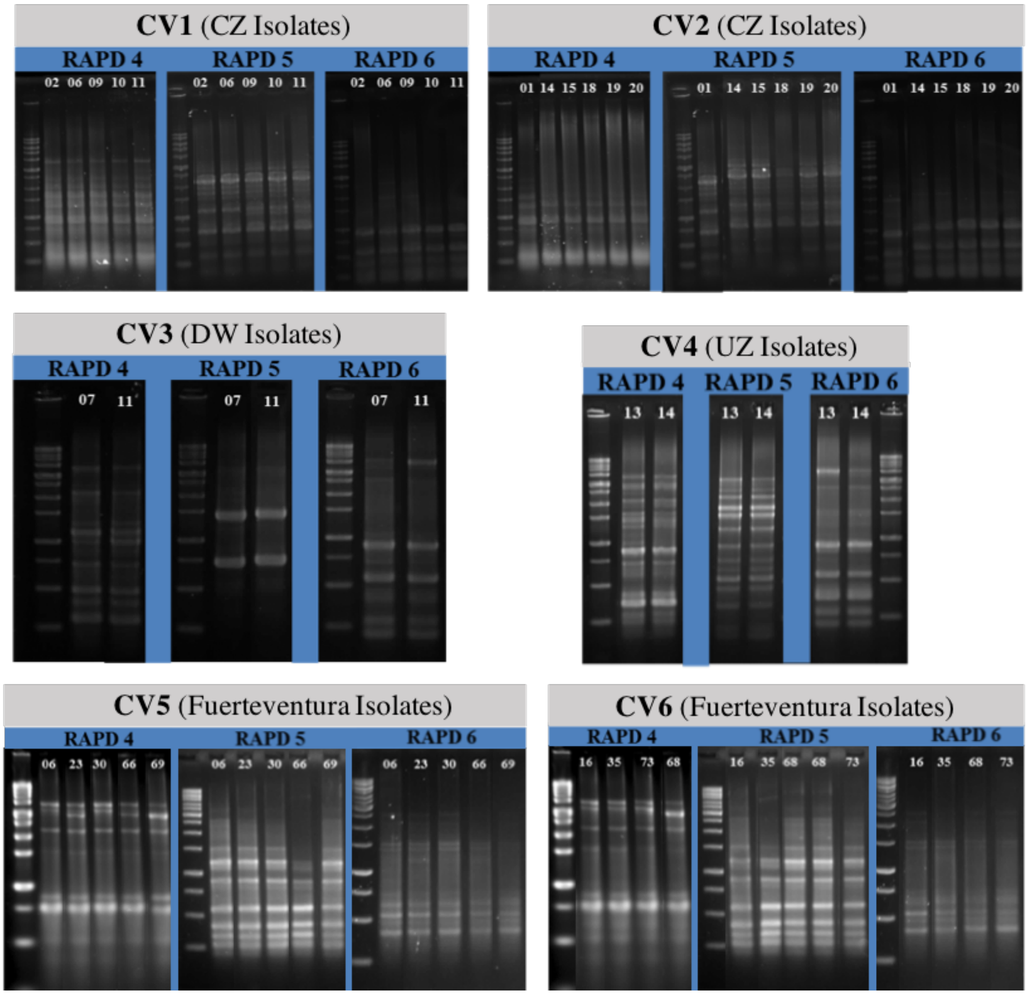

**Supplementary Figure S2:** Genomic diversity among *Sal. ruber* genomes recovered from the individual mesocosm experiments in the Mallorca solar salterns. The ANI value between pairs of genomes (x-axis) from the same mesocosm (graph title) is plotted against their shared genome fraction (y-axis). The graph to the top of each panel shows the number of ANI comparisons in each range of ANI value (0.1%). The mesocosm experiments were described previously (1, 2, 17).

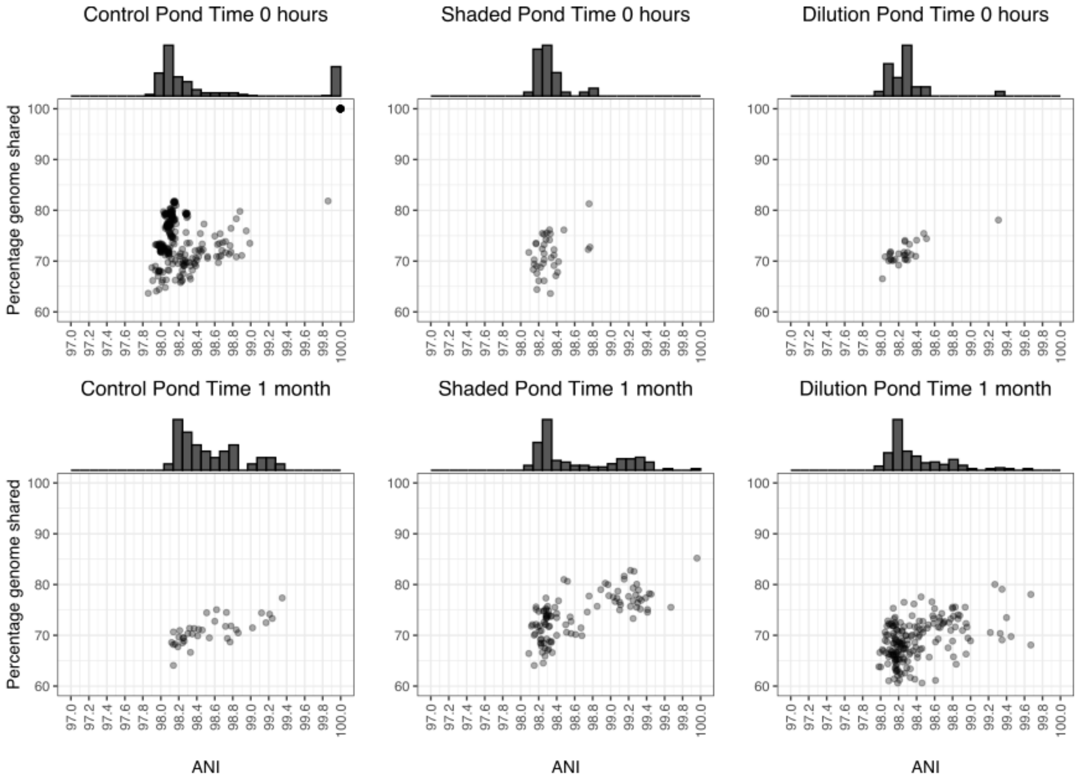

**Supplementary Figure S3:** Genomic diversity among *Sal. ruber* genomes recovered from the 'Fuerteventura' solar salterns. The ANI value between pairs of genomes (x-axis) is plotted against their shared genome fraction (y-axis). The graph to the top of shows the number of ANI comparisons in each range of ANI value (0.01%).

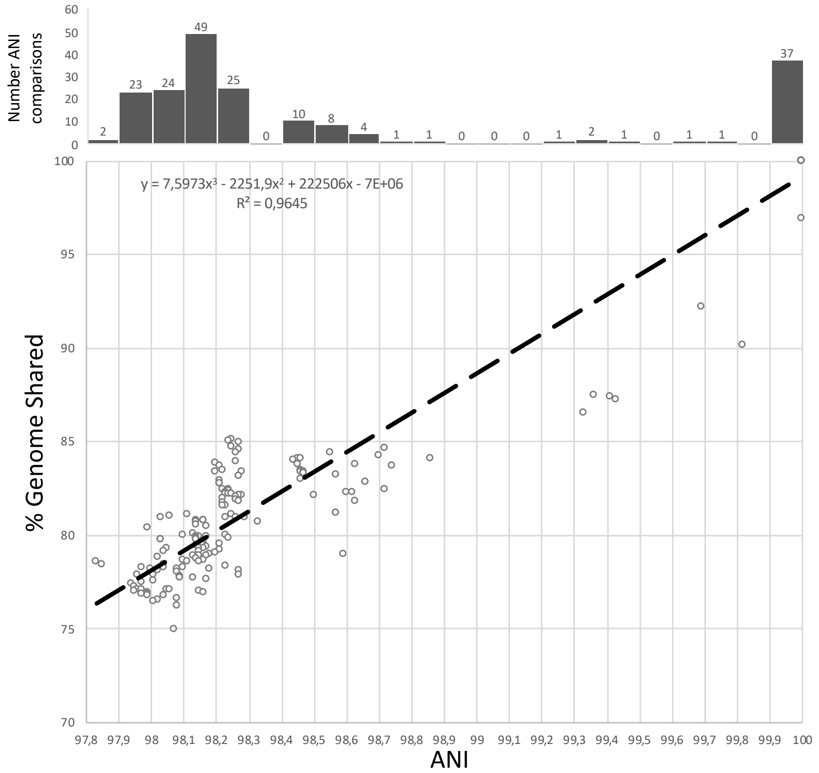

**Supplementary Figure S4:** Genomic diversity assessed by the number of different alleles within each core-orthology group (OG) identified. 793 OGs were assessed.

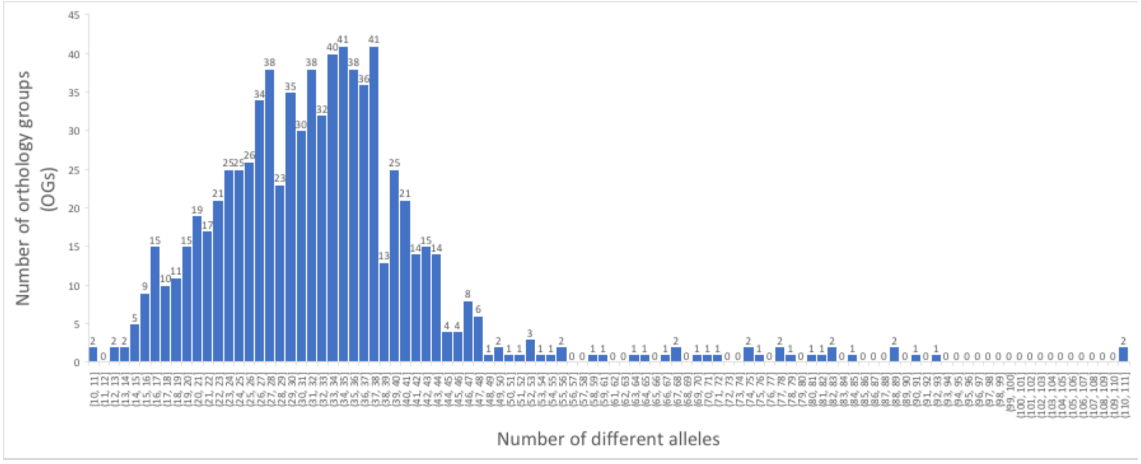

**Supplementary Figure S5:** Comparison between genomic clustering based on the percentage of alleles shared and phylogenetic analysis based on 793 core ortholog genes. To identify allelic variation in core genes among the genomes, the sequences assigned to a single OG were grouped using a 99.8% nucleotide sequence identity cut-off (see also Results section for further details). The percentage of alleles shared between genomes at this identity level was used as a distance metric to cluster the genomes, and the resulting dendrogram (on the left) is compared to a phylogenetic tree of the genomes using Neighbor Joining algorithm (on the right).

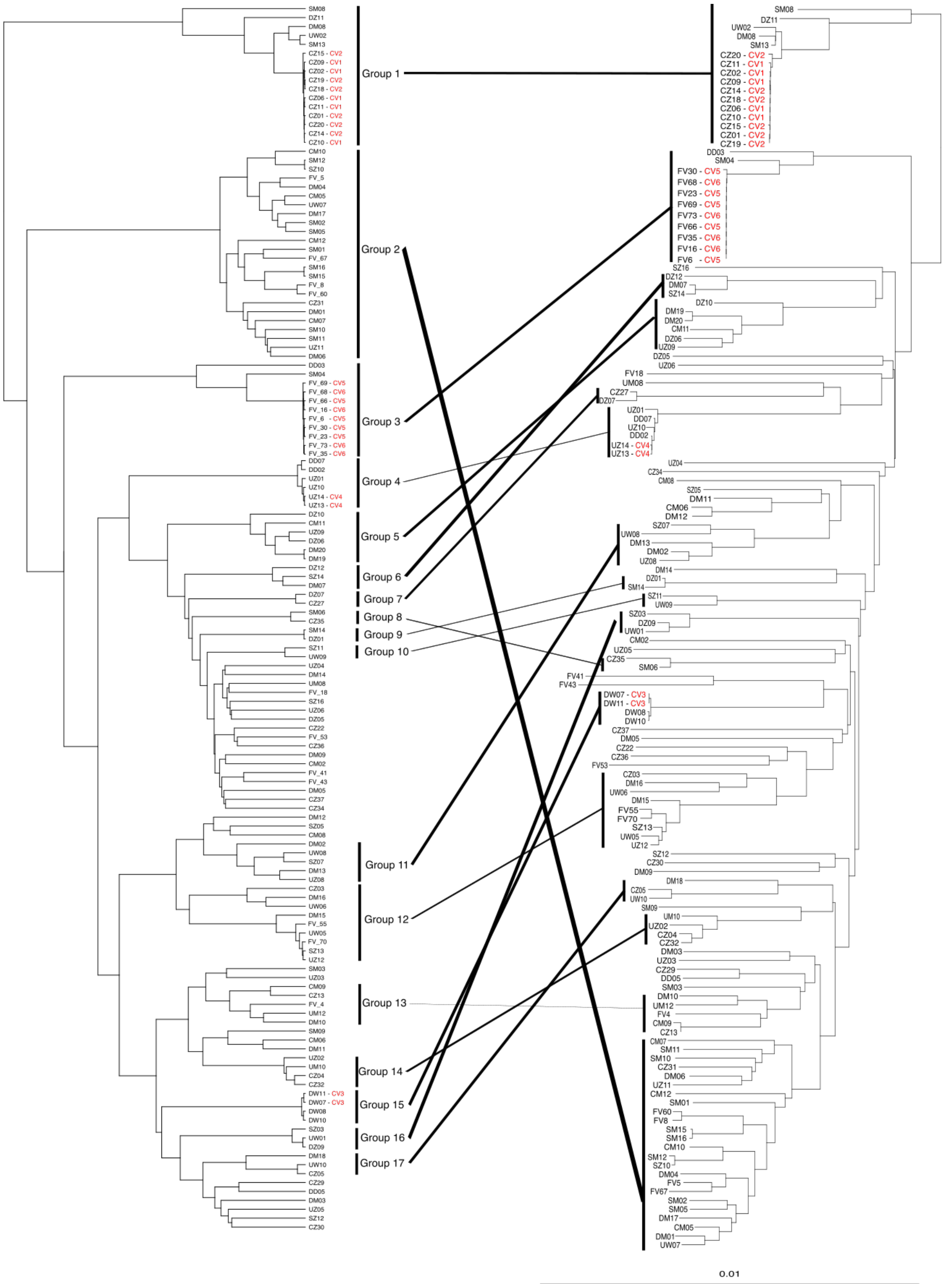

**Supplementary Figure S6: Genomic clustering of isolate genomes based on the percentage of alleles shared.** In the analysis, 793 core-OGs without paralogs (i.e., single-copy) were included. Alleles were defined as genes sharing at least 99.8% nucleotide identity. The percentage of alleles shared between genomes at this identity level was used as a distance metric to cluster the genomes, and the resulting overall similarity in gene-content was represented in a heatmap using the ggplot2 package v3.3.6 in Rstudio v1.1.4.

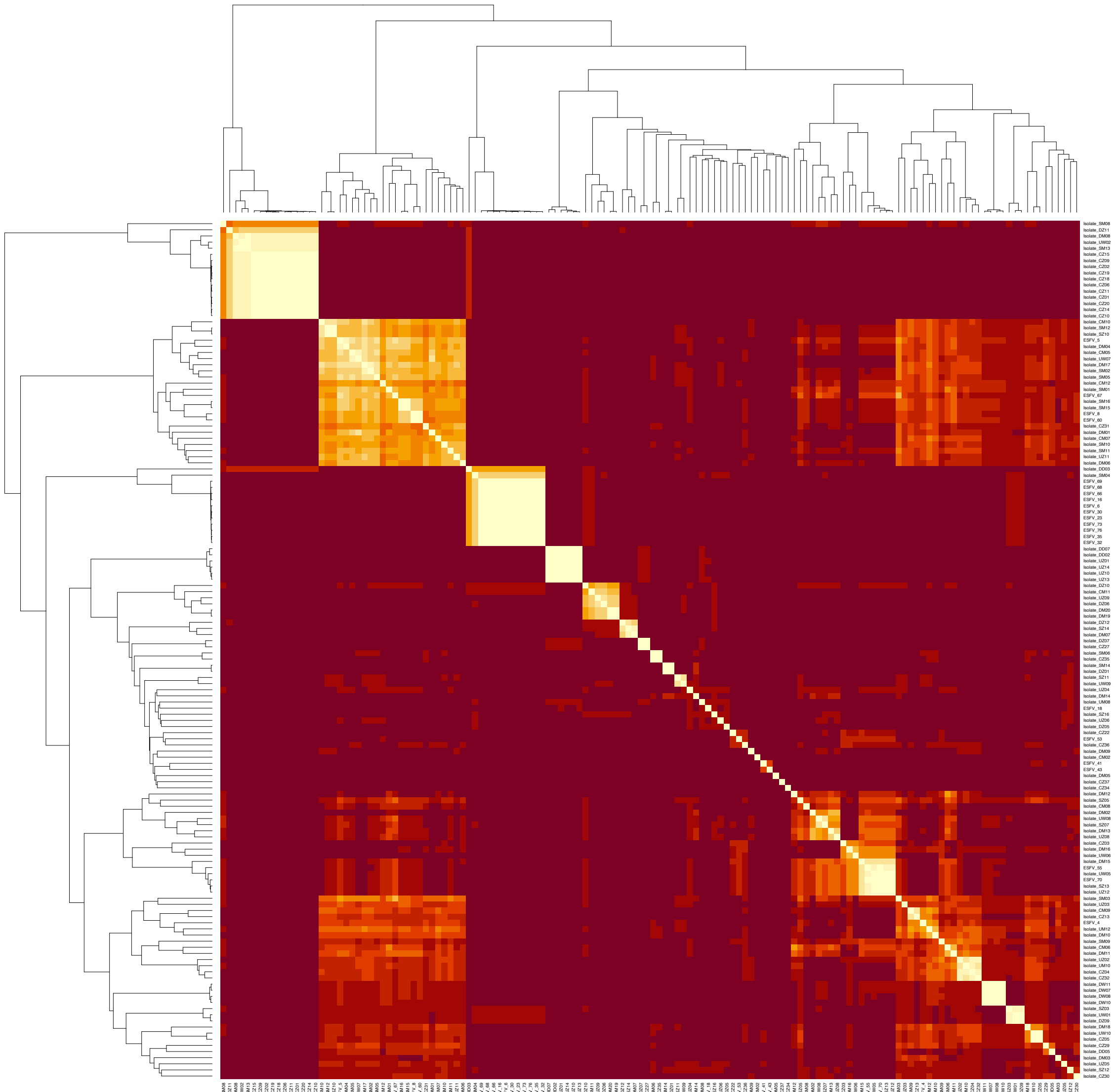

**Supplementary Figure S7: RAPD signatures of CV3 and comparison with their close relative isolates.** RAPD fingerprints were obtained using the RAPD4, RAPD5 and RAPD6 amplification primers (6). Each column in the gels shows a RAPD profile for a different isolate. Isolates DW07 and DW11 belong to CV3 are designated by blue fonts; closely related isolates are designated by other colors.

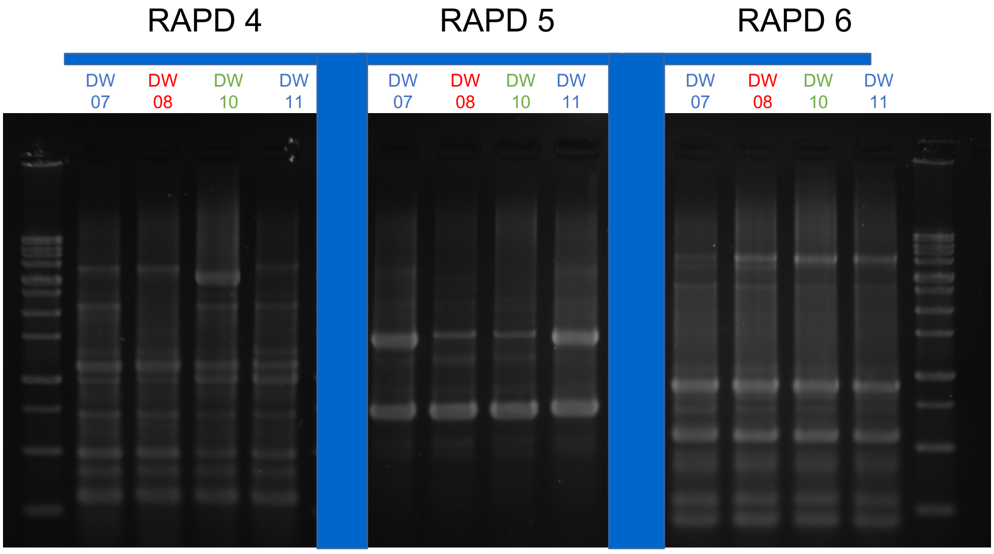

**Supplementary Table S1: Metagenomic relative abundance of the isolates representing different genomovars.** The table shows the relative abundance of the isolates based on metagenomic reads from the sample where the isolates were recovered. Relative abundance was estimated based on a competitive best-match approach to map reads to core genes as described in the main text. (A) Mallorca isolates (rows) and metagenomes (columns); (B) Fuerteventura isolates (rows) and metagenomes (columns).

**A)**

| Strains | Control Pond | | |
|---|---|---|---|
| | Control T0h | Control T1w | Control T1m |
| DM13 | 5.07 | 5.22 | 4.37 |
| UZ12 | 3.03 | 1.66 | 1.79 |
| CM09 | 2.89 | 3.07 | 3.05 |
| CZ13 | 2.86 | 3.00 | 3.42 |
| CZ31 | 1.74 | 1.66 | 1.87 |
| UZ02 | 2.76 | 3.46 | 3.47 |
| UZ08 | 2.62 | 2.72 | 2.56 |
| CZ27 | 2.45 | 2.95 | 2.72 |
| CM04 | 2.32 | 2.58 | 1.52 |
| UZ06 | 2.18 | 2.30 | 2.06 |
| CZ26 | 2.05 | 2.71 | 1.19 |
| CM08 | 1.94 | 2.42 | 2.58 |
| DM08 | 1.89 | 2.11 | 2.23 |
| DZ06 | 1.85 | 2.58 | 2.98 |
| UZ09 | 1.73 | 1.96 | 1.73 |
| CM06 | 1.57 | 1.65 | 1.48 |
| UZ05 | 1.57 | 1.69 | 1.64 |
| CZ29 | 1.57 | 1.79 | 1.80 |
| DM07 | 1.55 | 1.46 | 1.63 |
| CM05 | 1.54 | 1.70 | 1.79 |
| DM16 | 1.50 | 1.50 | 1.68 |
| DZ09 | 1.44 | 1.60 | 1.55 |
| CZ30 | 1.41 | 1.59 | 1.55 |
| UZ11 | 1.31 | 1.62 | 1.44 |
| DM02 | 1.23 | 1.04 | 1.17 |
| DZ07 | 1.20 | 1.27 | 1.47 |
| CZ05 | 1.19 | 1.48 | 1.58 |
| DZ10 | 1.18 | 0.94 | 1.15 |
| DM15 | 1.16 | 1.23 | 1.30 |
| UZ04 | 1.15 | 1.16 | 1.20 |
| DW07 | 1.10 | 1.50 | 1.78 |
| CZ22 | 1.01 | 1.05 | 1.18 |
| CZ03 | 1.00 | 0.89 | 1.23 |
| DM05 | 0.98 | 1.34 | 1.26 |
| UZ14 | 0.98 | 0.79 | 0.71 |
| CZ36 | 0.97 | 1.10 | 1.19 |
| CZ34 | 0.96 | 0.93 | 0.86 |
| DZ01 | 0.89 | 0.71 | 0.79 |
| UZ03 | 0.89 | 0.93 | 0.95 |
| M02 | 0.88 | 0.77 | 0.78 |
| CZ35 | 0.88 | 0.92 | 0.85 |
| DM03 | 0.85 | 0.79 | 0.76 |
| DM14 | 0.83 | 0.92 | 0.89 |
| DZ11 | 0.74 | 0.86 | 0.69 |
| CZ04 | 0.71 | 0.85 | 0.96 |
| CM11 | 0.69 | 0.50 | 0.47 |
| DM10 | 0.68 | 0.59 | 0.75 |
| UM10 | 0.65 | 0.87 | 0.87 |
| UM12 | 0.63 | 0.65 | 0.69 |
| DM01 | 0.63 | 0.55 | 0.60 |
| DM09 | 0.62 | 0.60 | 0.71 |
| DM19 | 0.60 | 0.52 | 0.71 |
| DZ12 | 0.58 | 0.65 | 0.64 |
| DZ04 | 0.56 | 0.59 | 0.44 |
| UM08 | 0.55 | 0.55 | 0.70 |
| DM18 | 0.51 | 0.57 | 0.59 |
| DM11 | 0.41 | 0.45 | 0.51 |
| CZ37 | 0.40 | 0.24 | 0.31 |
| DZ05 | 0.40 | 0.41 | 0.49 |
| CZ20 | 0.37 | 0.24 | 0.30 |

**B)**

| | Fuerteventura pond |
|---|---|
| FV6 | 10.62692386 |
| FV18 | 3.141455372 |
| FV41 | 18.31339059 |
| FV43 | 28.34565381 |
| FV4 | 4.667571254 |
| FV53 | 3.767981584 |
| FV55 | 5.65883573 |
| FV 5 | 2.778187827 |

**Supplementary Table S2:** Nonpareil analysis of the metagenomic reads representing the total *Sal. ruber* population in the control pond over a sampling period of one month.

| Nonpareil Statistics | Time zero | Time 1 week | Time 1 month | Concatenation of the three samples |
|---|---|---|---|---|
| **Redundancy** | 0.927 | 0.953 | 0.941 | 0.975 |
| **Average coverage** | 0.933 | 0.957 | 0.946 | 0.977 |
| **Actual sequencing effort** | 47,651,918 | 74,515,635 | 50,224,608 | 172,392,831 |
| **Pearson's coefficient** | 0.999 | 0.999 | 0.999 | 0.999 |
| **Estimated sequencing effort** | 45,957,814 | 40,539,528 | 39,110,605 | 39,934,906 |
| **Nonpareil sequence-diversity index** | 14.58 | 14.50 | 14.47 | 14.49 |

**Supplementary Table S3:** Estimations of the number of genomovars making up the total *Sal. ruber* population and 99% prediction interval (lower and upper boundaries).

| Sample (Control Pond) | Strains fit | Strains lower | Strains upper |
|---|---|---|---|
| **Time zero (CZT0h)** | 11,113 | 9,205 | 13,416 |
| **Time 1 week (CZT1W)** | 6,181 | 5,060 | 7,549 |
| **Time 1 month (CZT1M)** | 5,496 | 4,501 | 6,711 |
| **Concatenation of the three samples** | 6,775 | 5,605 | 8,190 |
